# *Naegleria fowleri:* protein structures to facilitate drug discovery for the deadly, pathogenic free-living amoeba

**DOI:** 10.1101/2020.10.20.327296

**Authors:** Kayleigh Barrett, Logan Tillery, Jenna Goldstein, Jared W. Lassner, Bram Osterhout, Nathan L. Tran, Lily Xu, Ryan M. Young, Justin Craig, Ian Chun, David M. Dranow, Jan Abendroth, Silvia L. Delker, Douglas R. Davies, Stephen J. Mayclin, Brandy Calhoun, Madison J. Bolejack, Bart Staker, Sandhya Subramanian, Isabelle Phan, Donald D. Lorimer, Peter J. Myler, Thomas E. Edwards, Dennis E. Kyle, Christopher A. Rice, James C. Morris, James W. Leahy, Roman Manetsch, Lynn K. Barrett, Craig L. Smith, Wesley C. Van Voorhis

**Affiliations:** Department of Medicine, Division Allergy and Infectious Disease, Center for Emerging and Re-emerging Infectious Disease (CERID) University of Washington, Seattle, WA, USA; Seattle Children’s Research Institute, Seattle, WA, USA; UCB Pharma, Bainbridge Island, WA, USA; Center for Tropical and Emerging Global Diseases, University of Georgia, Athens, GA, USA; Eukaryotic Pathogens Innovation Center, Department of Genetics and Biochemistry, Clemson University, Clemson, SC, USA; Department of Chemistry, University of South Florida, Tampa, FL, USA; Department of Chemistry and Chemical Biology and Department of Pharmaceutical Sciences, Northeastern University, Boston, MA, USA; Seattle Structural Genomics Center for Infectious Diseases, Seattle, WA, USA; Department of Biology, Washington University, St. Louis, MO, USA; Department of Pediatrics, University of Washington, Seattle, WA USA; Department of Biomedical Informatics & Medical Education, University of Washington, Seattle, WA, USA; Department of Global Health (Pathobiology), University of Washington, Seattle, WA, USA

**Author notes:** These authors contributed equally to this work (Jenna Goldstein, Jared W. Lassner, Bram Osterhout, Nathan L. Tran, Lily Xu, Ryan M. Young). Department of Pharmaceutical and Biomedical Sciences, College of Pharmacy, University of Georgia, Athens, Georgia, USA.

## Abstract

*Naegleria fowleri* is a pathogenic, thermophilic, free-living amoeba which causes primary amebic meningoencephalitis (PAM). Penetrating the olfactory mucosa, the brain-eating amoeba travels along the olfactory nerves, burrowing through the cribriform plate to its destination: the brain’s frontal lobes. The amoeba thrives in warm, freshwater environments, with peak infection rates in the summer months and has a mortality rate of approximately 97%. A major contributor to the pathogen’s high mortality is the lack of sensitivity of *N. fowleri* to current drug therapies, even in the face of combination-drug therapy. To enable rational drug discovery and design efforts we have pursued protein production and crystallography-based structure determination efforts for likely drug targets from *N. fowleri. N. fowleri* genes were selected if they had homology to drug targets listed in Drug Bank or were nominated by primary investigators engaged in *N. fowleri* research. In 2017, 178 *N. fowleri* protein targets were queued to the Seattle Structural Genomics Center of Infectious Disease (SSGCID) pipeline, and to date 89 soluble recombinant proteins and 19 unique target structures have been produced. Many of the new protein structures are potential drug targets and contain structural differences compared to their human homologs, which could allow for the development of pathogen-specific inhibitors. Five of the structures were analyzed in more detail, and four of five show promise that selective inhibitors of the active site could be found. The 19 solved crystal structures build a foundation for future work in combating this devastating disease by encouraging further investigation to stimulate drug discovery for this neglected pathogen.

## 1. INTRODUCTION

In Australia in 1965, Fowler and Carter reported the first case of *Naegleria fowleri* infection, commonly referred to as the “brain-eating amoeba”, which is the only known species of the *Naegleria* genus associated with human disease (1). The free-living amoeba is found in soil and freshwater on all seven continents, but is mainly found in warmer regions flourishing in freshwater and soils with higher temperatures up to 115°F (46°C) (2).

Infection rates of the amoeba increase during summer months causing the disease, primary amebic meningoencephalitis (PAM) (2, 3). PAM is a fatal CNS disease that displays severe meningitis and cranial pressure caused by inflammation of the brain (3, 4). The National Institute of Allergy and Infectious Diseases (NIAID) has classified *N. fowleri* as a category B priority pathogen, the second highest class of priority organisms/biological agents. Category B pathogens typically have high mortality rates, are easily disseminated, may cause public panic and social disruption, and require special action for public health preparedness (2, 5).

Between the years 1962 and 2016, 145 cases were reported by the CDC within the USA, and only 4 patients (3%) survived *N fowleri* infection and subsequent PAM (6). This 54-year reporting tally might suggest only several PAM cases occur in the USA a year. However, a recent review found that hundreds of undiagnosed cases of fatal “meningitis and encephalitis” were reported in the summers between the years 1999 and 2010 in persons aged 2-22 years in the Southeast USA (7). Cases lacking diagnostic brain biopsy or spinal fluid analysis of PAM may account for a portion of the undiagnosed, inconclusive cases (7). Thus, the incidence of *N fowleri* infections in the USA is probably much higher than several a year arguing, that *N. fowleri* poses a much higher public health threat than the documented cases would suggest (7). Rising global temperatures has also led to claims of increased risk of infections due to spread of suitable water conditions for *N. fowleri* (3, 8).

The global burden of *N. fowleri* PAM cases is likely greater than anticipated, yet undefined. Aga Khan University in Karachi, Pakistan has reported a rise in the number of PAM cases. Aga Khan Hospital serves only a small fraction of the Karachi population (∼23 million total) and reports around 20 cases/year of PAM. Other hospitals in Karachi did not report a single case during the same time, likely due to the lack of awareness, autopsies, and microscopy (9). Only about 10 cases of *N. fowleri* PAM have been reported in Africa, though the numbers are likely impacted by a reporting bias, as the resources for diagnostic brain biopsy are rarely present, and autopsies are almost never performed (10).

The combination of a devastating mortality rate, a warming climate, and a rapid-onset infection emphasize why *N. fowleri* should not stay a neglected organism. Treatment is extremely limited and not well defined, using a combination of known antifungals, antibiotics and microbicides (11). Amphotericin B is the current drug of choice, in high dosages, in combination with other repurposed drugs such as rifampin, miltefosine, and fluconazole (9, 12). Recent studies suggest that posaconazole is more efficacious than fluconazole in vitro and in animal models of PAM (12). Miltefosine was used in combination to successfully treat two patients, one reported and one unreported in the literature. Miltefosine in combination is not always helpful, in that a patient was treated with miltefosine and suffered permanent brain damage and another had a fatal outcome (11, 13). The multi-drug therapy is associated with severe adverse effects and requires higher than normal dosages to penetrate the blood-brain barrier and to reach the CNS (11, 14). New development of rapid-onset, brain permeable, efficient, and safe drugs is urgently needed.

Given the lack of understanding about causes of drug resistance of *N. fowleri* and the urgent need for new drugs, we investigated the proteome of *N. fowleri* for likely drug targets attempting to enable further drug discovery efforts by producing material for characterization of the proteins. This work is a first step towards the discovery of drugs specifically designed against *N fowleri*.

## 2. RESULTS

### 2.1. The *N fowleri* proteome contains hundreds of potential drug targets

Potential drug targets were selected by sequence homology to DrugBank protein targets (15). Additional targets were requested by the amoeba research community, leading to a total of 178 *N. fowleri* targets entering the Seattle Structural Genomics for Infectious Disease (SSGCID) structure determination pipeline. The SSGCID is a National Institutes for Allergy and Infectious Disease (NIAID) supported preclinical service for external investigators (www.SSGCID.org) (15). All targets were filtered according to the standard SSGCID target selection protocol and criteria (13): eliminating proteins with over 750 amino acids, 10 or more cysteines, or 95% sequence identity with 70% coverage to proteins already in the PDB, targets claimed or worked on by other scientific groups, and targets with transmembrane domains (except where a soluble domain could be expressed separately) (18). Target criteria resulted in selection of 178 proteins which entered the SSGCID production pipeline. These proteins, homologous to other known drug targets, consisted of metabolic enzymes, protein synthetases, kinases, and others. Real-time updates to target status progress can be viewed at the SSGCID website. Figure 1 shows a view of current status. Additionally, a detailed table of the *Naegleria* protein crystallography statistics is available in Supplemental Information.

**Figure 1.**
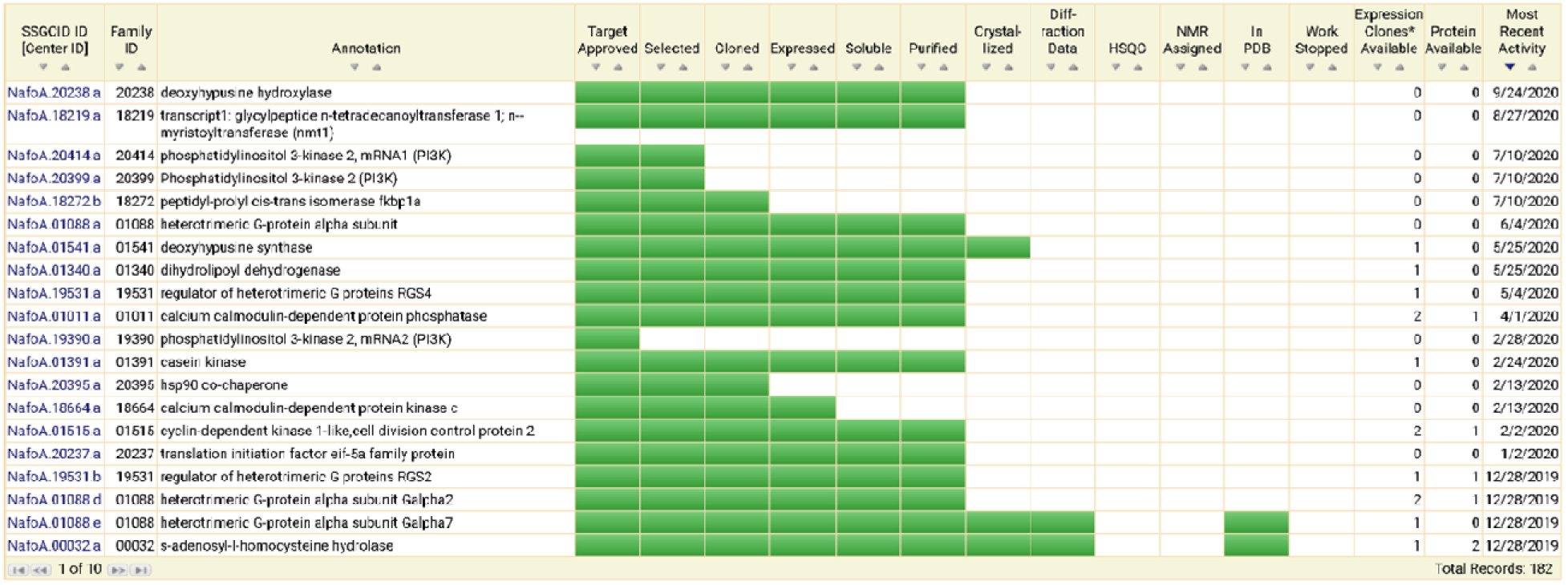
Status of *Naegleria* targets in the SSGCID pipeline as of 10/1/2020. This view is publicly available at www.ssgcid.org through the Targets tab, sorting by genus. Targets are displayed by SSGCID identifier and annotation with current status shown by green bars, sorted here by most recent activity reported to the database. The number of expression clones is shown and indicates different forms of the target that are available to external users and can be ordered. Protein availability also indicates samples of protein available to the external community and represents at least one vial of concentrated protein of approximately 100 microliters ranging from 10-50 mg/mL of >95% purity material.

### 2.2. One third of targets attempted produced soluble protein

The open reading frames of each target were obtained from AmoebaDB.org. Progression of the targets through the SSGCID protein production pipeline is shown in Table 1. Of the 178 NIAID approved targets, 177 were selected for cloning. One protein target was eliminated due to redundancy given its 100% identity match and 84% coverage to another protein target already in the SSGCID pipeline. Of the 177 targets attempted in PCR amplification using *N. fowleri* cDNA, 133 were successfully amplified and cloned into SSGCID expression vectors (75%) (17). In small-scale expression screening, 82 of the 133 successfully cloned targets (61.6%) demonstrated soluble expression with a N-terminal His6-tag vector (17). Of these 82 soluble proteins, 64 proteins were purified to >95% purity with yields ranging from 1.1 mg to 348 mg. These proteins are available, under request, at SSGCID.org.

**Table 1.** Target status of *N fowleri* proteins within SSGCID pipeline.

|  |  |
| --- | --- |
| Target approved | 178 |
| Selected | 177 |
| Cloned | 133 |
| Expressed | 98 |
| Soluble | 82 |
| Purified | 64 |
| Crystallized | 29 |
| Diffraction | 27 |
| Native diffraction data | 19 |
| In PDB (apo) | 19 |

### 2.3. *N. fowleri* proteins resulted in a 11% structure determination success rate

Following purification of high-purity preparations of protein, the targets were submitted for structure determination by X-ray crystallography. Of the 64 proteins produced, 26 crystallized (29%), and 20 diffracted, 19 of which met the SSGCID resolution quality criteria and were submitted to the Protein Data Bank (PDB). Table 2 lists the 19 targets deposited in PDB that will be reported in this paper, including five structures with a unique ligand bound, for a total of 23 PDB deposits. Overall, this resulted in a structure determination success rate of 11%, which is comparatively higher than usual structural genomic pipeline rates reported by us or other structural genomic groups (19).

**Table 2.**
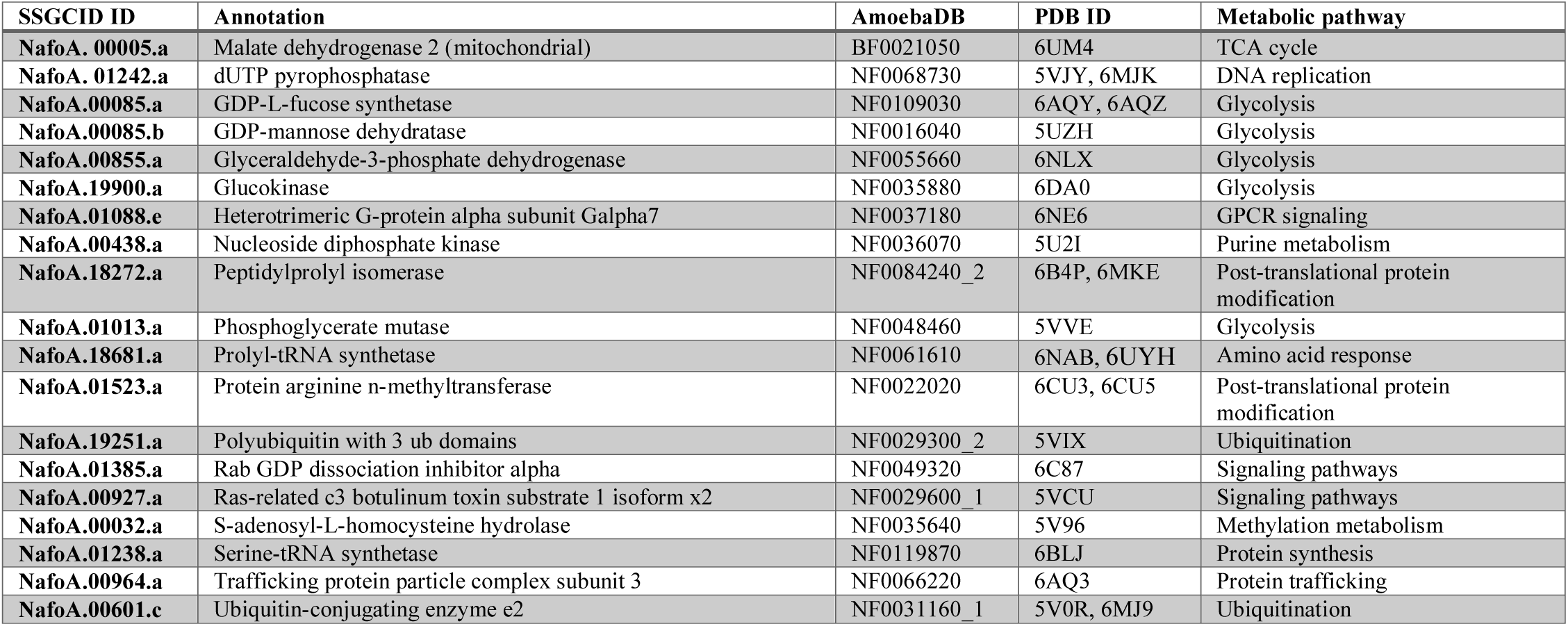
Listing of *Naegleria fowleri* structures deposited by SSGCID in the PDB.

### 2.4 Comparative analysis of *N. fowleri* and *Homo sapiens* enzyme active sites suggest some targets have promise for selective inhibition

We have already published that glucokinase inhibitors can be obtained that are selective for *N. fowleri* vs. the human homolog, supporting glucokinase as a target for *N. fowleri* therapeutics (20). We analyzed the other structures determined, comparing the *N. fowleri* structure to human homolog structures, in order to determine opportunities for selective design of chemical inhibitors. Comparison of the *N. fowleri* determined structure to human structures available in the PDB was done by superimposition of the coordinate files (Table 3). With the exception of a pair of 96% identical *N. fowleri* and human ubiquitin-conjugating enzymes e2, all of the *N. fowleri* enzymes differed from human homologs by more than 38% (Table 3). We wanted to focus on the known ligand binding sites, to search for potential differences for inhibitors. A PDB search revealed that five of the 19 *N fowleri* structures determined also had human homolog structures determined which contained a known inhibitor of the human protein (Table 3). We then manually inspected and compared the binding sites of these five proteins, described below for each protein. Despite sequence similarities of the active sites, there were four cases where a case for active site specificity could be made, supporting these proteins as targets for therapeutics for *N. fowleri*.

**Table 3.**
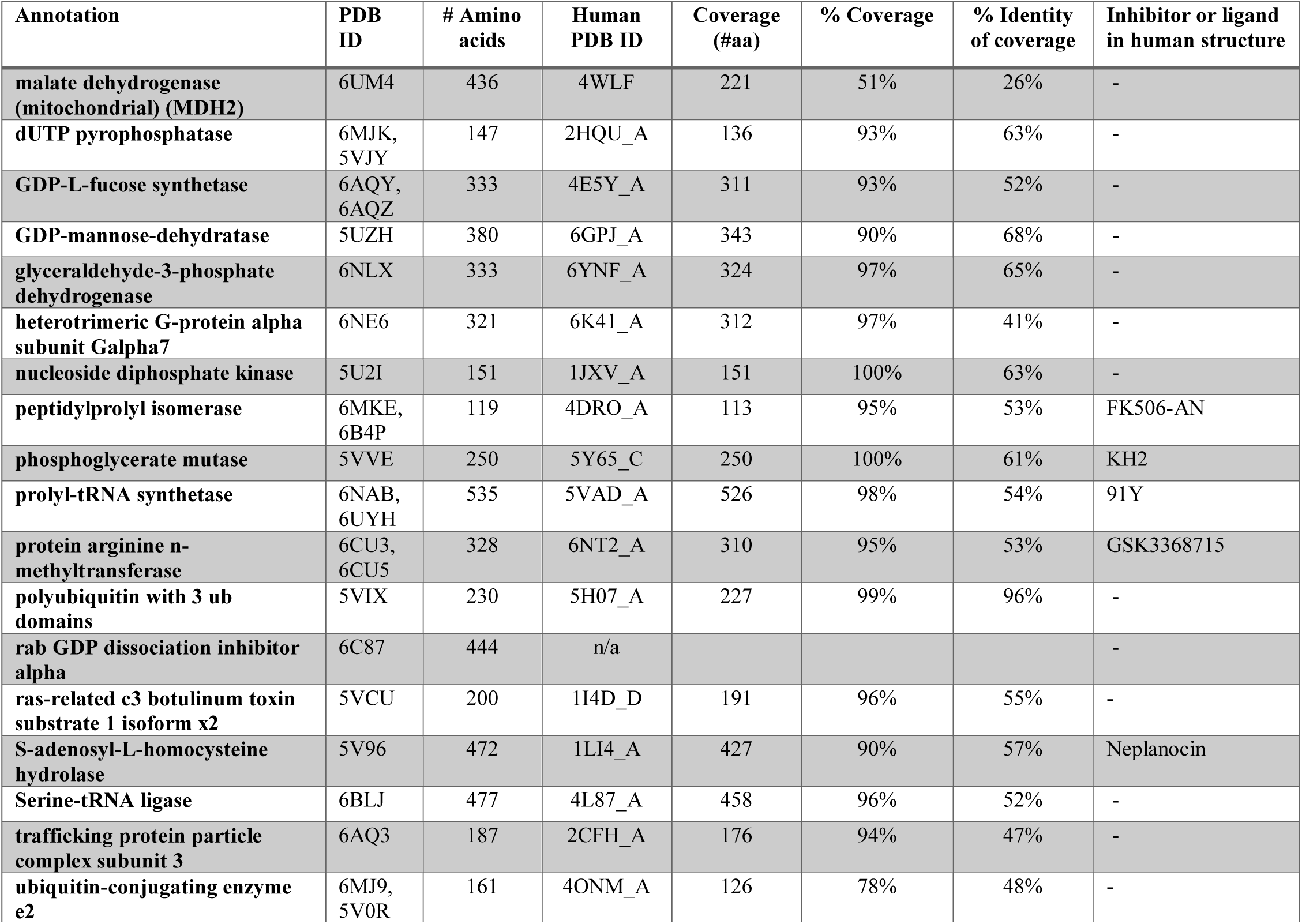
Comparative analysis of *N fowleri* and human structures deposited the PDB.

*N. fowleri* S-adenosyl-L-homocysteine hydrolase (*Nf*SAHH) catalyzes the breakdown of S-adenosyl-homocysteine (SAH) into adenosine and homocysteine. SAH is a byproduct of S- adenosyl-L-methionine as a methyltransferase; the transfer of a methyl group to its respective cellular substrates such as DNA or rRNA, produces SAH (21). SAH hydrolases play a central role in methylation reactions required for growth and gene regulation, and inhibitors of SAH hydrolase are expected to be antimicrobial drugs, especially for eukaryotic parasites (21). Ribavirin is structurally similar to adenosine and has been proved to produce a time-dependent inactivation of human *(Hs)* SAHH and *Trypanosoma cruzi* (*Tc*) SAHH (22).

The *Nf*SAHH asymmetric unit contains a homo-tetramer (Figure 2). Although each chain contains an active site, structural analysis indicates that two chains must be present for the hydrolysis reaction to occur successfully. Each chain consists of three domains: a substrate- binding, a cofactor-binding, and a C-terminus domain (23). When substrates are not bound, the substrate-binding domain is located on the exterior, far from the meeting point of all four subunits of the asymmetric unit (24). The C-terminus domain is involved in both cofactor binding and protein oligomerization (23). In addition to the three main constituents, the structure contains two hinge regions that connect the substrate-binding and cofactor-binding domains. When substrates bind, the hinge region changes conformation, closing the cleft between the substrate-binding domain and the cofactor-binding domain of the respective chain (24). In the structure of MSAHH, all subunits exhibit a closed conformation.

**Figure 2.**
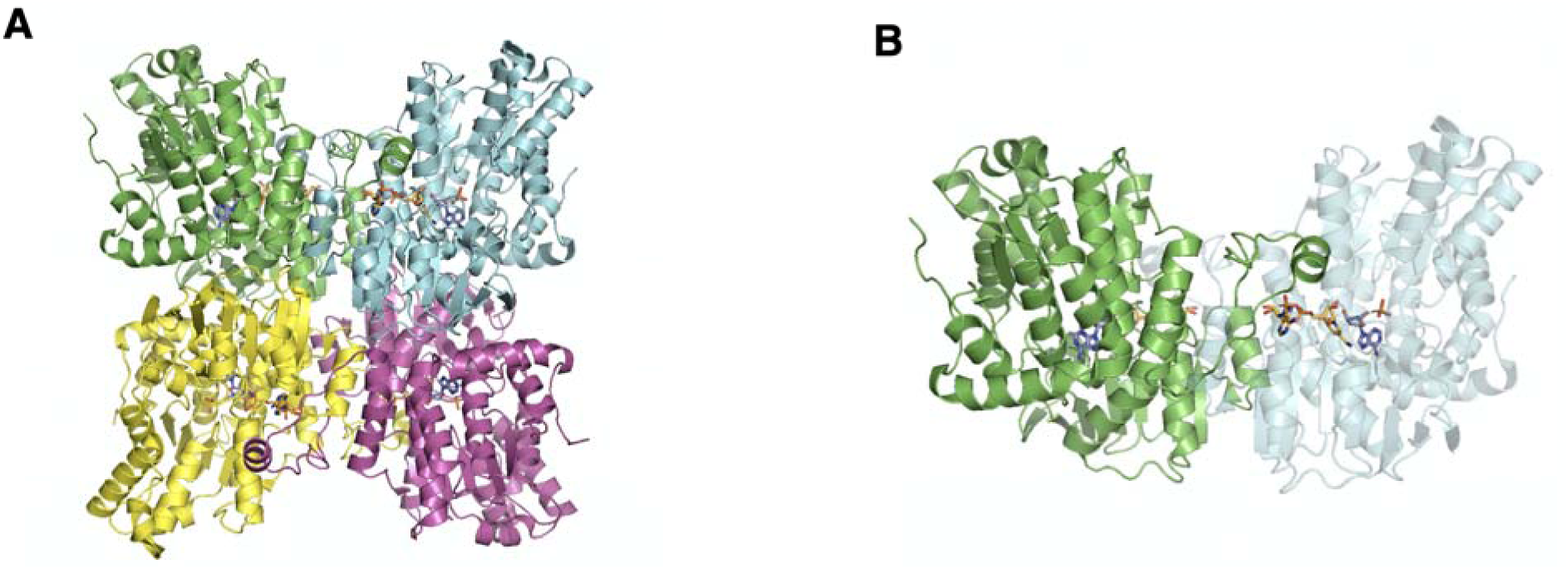
Crystal structure of *Nf*SAHH (PDB: 5V96) solved at a 2 Å resolution. (**A**) The asymmetric unit of SAHH. Individual polypeptide chains are colored green (chain A), pink (chain B), yellow (chain C), and cyan (chain D). (**B**) The biological unit is a homotetramer with a 2-fold axis of symmetry. Each of the four chains has its own active site containing one NAD^+^ molecule (yellow), one adenosine molecule (purple), and a phosphate (orange).

SAHH is one of the most highly conserved proteins among species, with many of the same amino acids binding the same substrates across homologs. *Nf*SAHH is 62% identical to the human homolog. In the NAD binding region, conserved Lys and Tyr bind via hydrogen bonds to oxygens of NAD in both *Nf*SAHH and *Hs*SAHH (24) (Figure 3). Residues involved with binding adenosine (ADO) are also highly conserved, with Gly341-His342-Phe333 being completely conserved (24). To regulate the entrance of substrates into/out of the active site, there is a highly conserved His-Phe sequence within the cofactor-binding domain. This works as a molecular gate that, when the protein is in open conformation, allows access to the substrate-pocket (23).

**Figure 3.**
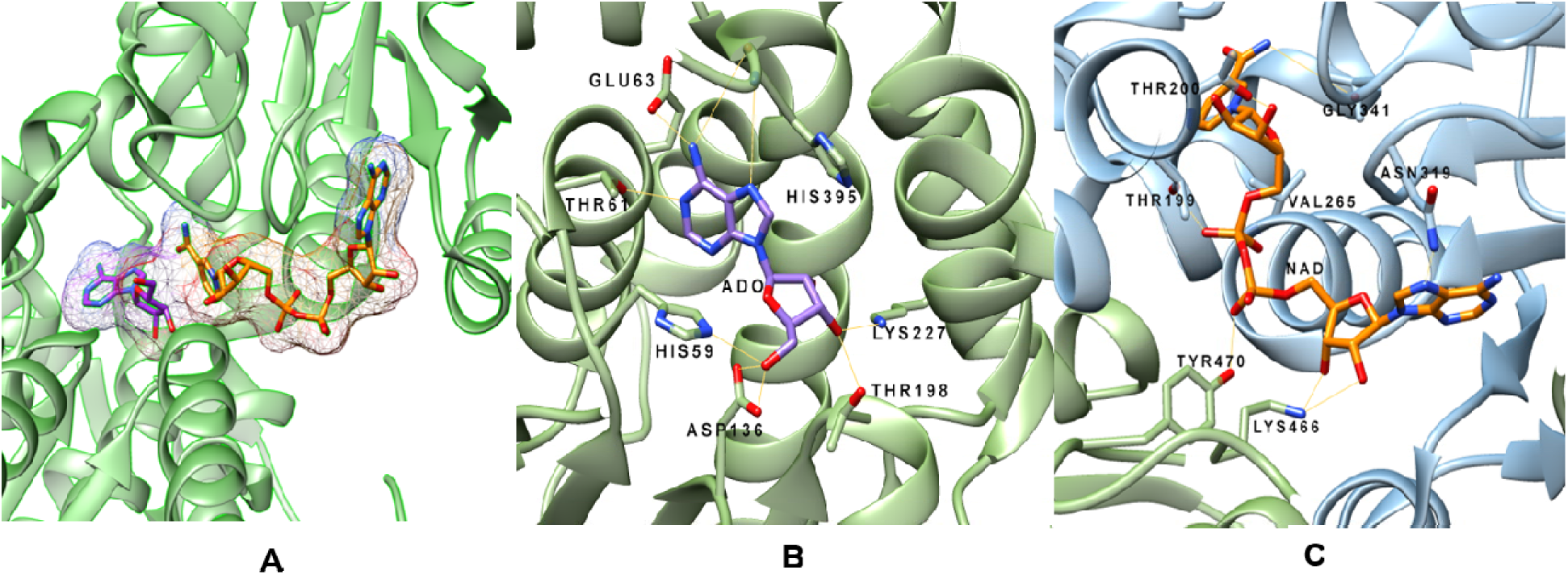
Overview of NfSAHH active site binding. (**A**) Nicotinamide adenine dinucleotide (NAD/orange sticks) and adenosine (ADO/purple sticks) both fit well within the structure of SAHH. (**B**) Hydrogen-bond network denoted by yellow dashed lines around ADO involving chain A. (**C**) Hydrogen-bond network involving NAD with chain A (blue) and chain B (green).

A search of the PDB for *Hs*SAHH structures found multiple inhibitor bound human structures including 1LI4 (neplanocin) and 5W49 (oxadiazole compound). A comparison of the *Nf*SAHH structure to the neplanocin bound human structure revealed a highly conserved conformation of the protein. In the neplanocin bound structure, the two domains of the monomer are similar to the *Nf*SAHH structure. However, in the oxadiazole bound structure, two domains of the monomer are in a more open conformation, where the C-terminal and N-terminal domains have opened up relative to each other in a hinge-opening motion. The oxadiazole compound stretches across the interface and is surrounded by 11 residues within 4 Å. Of the 11 residues coordinating the inhibitor oxadiazole compound of the 5W49 structure, 10 are identical between the *Naegleria* and human SAHH. Only one change of M351T relative to the human enzyme is present, suggesting a highly conserved inhibitor binding site.

The crystal structure of *Nf*SAHH (Figure 4) contains the adenosine substrate and NAD cofactors bound to the active site to guide structure-activity relationships that could help to optimize adenosine analog compounds. The sequence differences that line the access channel at the dimer interface allow a rational approach to selectively inhibit the otherwise highly conserved active site (25). Amoeba SAHHs have an additional helix insertion that in *Nf*SAHH forms a hydrophobic groove accessible from the adenosine binding site (Figure 4 B, C). Specificity could be achieved by designing compounds that simultaneously target this hydrophobic pocket and the active site (Figure 4). Thus, we feel a reasonable case can be made that structural differences, close to the active site, would allow development of specific *Ng*SAHH inhibitors supporting development of a therapeutic.

**Figure 4.**
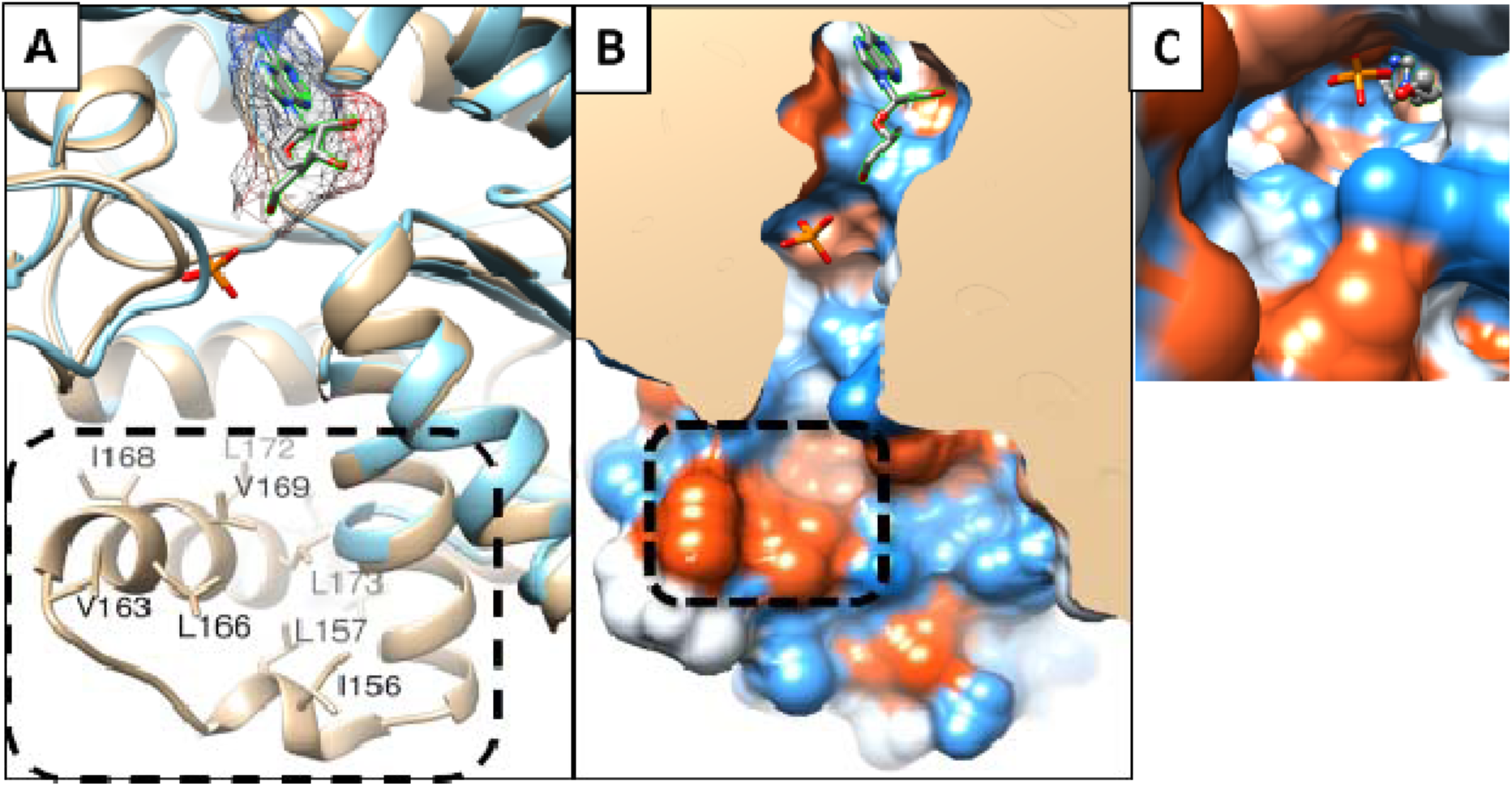
SAHH active site analysis. (**A**) Adenosine-bound *Nf*SAHH (PDB: 5V96, tan) vs. *Hs*SAHH bound to adenosine analogue (PDB: 1A7A, blue); box highlights *Nf*-specific insertion with labelled hydrophobic residues. (**B**) Slice through *Nf*SAHH, with surface colored on Kite-Doolittle (blue-red) hydrophobicity scale; dashed box indicates hydrophobic groove formed by Val163, Leu166, Val169 and Leu173 at opening of deep-seated adenosine pocket. (**C**) Same view as (**B**) tilted 90° shows opening from hydrophobic groove to adenosine pocket.

***N. fowleri*** **phosphoglycerate mutase (*Nf*PGM)**, a glycolysis enzyme, catalyzes the isomerization of 3-phosphoglycerate and 2-phosphoglycerate during glycolysis and gluconeogenesis and is regarded as a key enzyme in most organism’s central metabolism (26). There are two distinct forms of PGMs, differentiated by their need of 2, 3-bisphosphoglycerate as a cofactor. PGM in mammals require the cofactor whereas PGM present in nematodes and bacteria do not (27). The MPGM is likely the cofactor-dependent PGM type. The crystal structure of MPGM (PDB: 5VVE) was solved at a resolution of 1.7 and consists of 250 amino acid residues (∼30 kDa).

The *Hs*PGM and *Nf*PGM structures share 61% identity. Residues surrounding the binding pocket for citrate acid are all conserved, with the exception of a conservative change from a Thr30 *(Nf)* to Ser30 *(Hs).* A comparison of the *Nf*PGM structure to the homologous human enzyme *Hs*PGM (PDB: 5Y65) shows a conformational opening of the substrate binding site to accommodate the KH2 ligand. However, the residues surrounding the inhibitor molecule and supporting the movement of the peptide are identical between the two enzymes. It is likely that with this high homologous identity that *Nf*PGM is not a strong candidate for selective active site inhibitor design.

***N. fowleri*** **protein arginine N-methyltransferase (*Nf*PRMT1)** methylates the nitrogen atoms found on guanidinium side chains of arginine residues within proteins. The methylation of nucleotide bases is a well-known mechanism of importance that influences DNA, nucleosomes, and transcription functionalities (28). The enzyme is highly conserved across eukaryotes. Faulty regulation or deviating expression of PRMTs is associated with various diseases including inflammatory, virus-related, pulmonary, and carcinogenesis (29). Overexpression of PRMTs has been observed in multiple forms and types of cancer, including PRMT1v1 overexpression in colon cancer (30) and large increases of PRMT1v2 in breast cancer (31). Inhibitor discovery and testing using PRMTs in cancer has been frequently employed (29). *Nf*PRMT1 was compared to the drug bound structure of *Hs*PRMT1 (PDB: 6NT2). The protein binds ligands at a dimer interface closing around two inhibitor molecules, one on each monomer. A large ligand binding loop is disordered in the *Nf*PRMT1 structure, presumably becoming ordered and visible in the crystal structure in the presence of inhibitor in the human structure. Due to the large binding surface for peptide substrates, PRMTs typically are promiscuous in nature with a wide range of binding substrates (29). Comparison of over 40 PRMT-inhibitor complexes revealed 5 distinct binding mechanisms at multiple sites including active site and allosteric binding pockets (32). Isozyme specific peptide mimics have been identified which preferentially bind *Hs*PRMT1 vs. *Hs*PRMT5 enzyme. A similar approach could be considered for selective *Nf*PRMT inhibitor development (33, 34). There is still a need to improve both the affinity and selectivity of these micromolar, sub-micromolar potent PRMT inhibitors as well as to better understand the enzyme’s biological and disease processes in greater scope (35).

Despite high sequence identity in the ligand binding pocket, there are distinct differences in side chain orientation between the structures. These residues may change conformation upon binding inhibitor. A number of distinct features of *Nf*PRMT1 exist which can be exploited for potential structure-based approach to developing selective allosteric inhibitors against the *Nf* enzyme. A methionine is present in the *Nf*PRMT1 structure adjacent to the adenine moiety of the S-adenosyl homocysteine (SAH) which differs significantly from all nine-known human PRMTs. The substrate binding region is lined by residues variant between *Nf* and all nine-known human PRMTs; for example, though the *Nf*PRMT1 pocket is similar to the allosteric inhibition pocket of *Hs*PRMT3, there are two tyrosine substitutions lining the pocket (36). Additionally, N-terminal residues which interact with inhibitors of *Hs*PRMT1 are largely not present or have limited interactions in *Nf*PRMT1 (37). Thus, inhibitors that selectively target *Nf*PRMT1 vs. the 9 *Hs*PRMTs are envisioned due to structural differences near the ligand binding sites.

***N. fowleri*****peptidylprolyl isomerase (*Nf*PPI)** is a member of a superfamily of proteins comprised of three structurally distinct main families: cyclophilins, FK506 binding proteins (FKBPs), and parvulins. Based on structural and sequence alignment, the *N. fowleri* structure falls in the FKBP family, a group of enzymes inhibited by compounds such as FK506 and rapamycin (38). PPIs assist protein folding and influence protein denaturation kinetics by catalyzing the cis/trans isomerization of peptide bonds preceding prolyl residues (39). The enzymes participate in a diverse array of processes ranging from signal transduction to gene regulation and have been found to have close interaction with heat shock 90 proteins (40). PPI inhibitors are an emerging class of drugs for many therapeutic areas including infectious diseases and many potent small molecule inhibitors have been derived for each of the members of the superfamily. However, selective inhibitor design has been difficult due to the shallow, broad, solvent-exposed active sites and their conservation between homologs and protein families (41).

The interior of the binding pocket of *Nf*PPI is mostly hydrophobic (Figure 5). Only four putative hydrogen-bonding interactions are observed between the enzyme and substrate. All residues involved in polar interactions in the NfPPI are also present in the human homolog *Hs*FKBP51 (PDB: 1KT0), but the regions occupied by two hydrophilic residues in *Hs*FKBP51(Ser118 and Lys121) are instead occupied by hydrophobic residues, (Ile and Leu, respectively) (40). Another difference found in the conformation of this loop region is the insertion of an additional residue after Gly95 of *Nf*PPI. These changes in structure and sequence may lead to selective inhibition and thus establish PPIs as a selective drug target for *Naegleria.*

**Figure 5.**
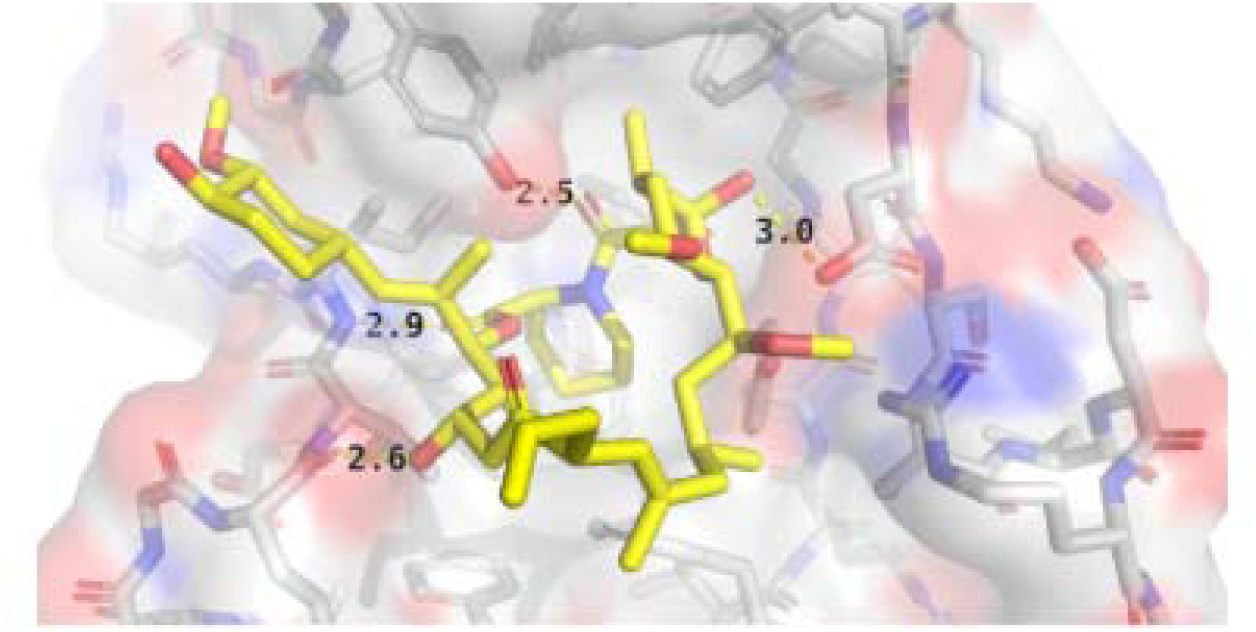
Interactions between *NfPPI* and FK506. FK506 (yellow-carbon stick model) sits inside a mostly hydrophobic binding pocket of *NfPPI* (white-carbon stick and surface model) consisting of Tyr38, Phe58, Trp71, and Phe111 on the distal side of the substrate. Hydrogen-bonding interactions exist between FK506 and the side chains of Asp49 and Tyr94, and the backbones of Glu66 and Ile68.

***N. fowleri*** **Prolyl-tRNA synthetase (NfProRS)**. Aminoacyl-tRNA synthetases (ARSs) are globally essential enzymes among all living species. Their roles in protein translation and biosynthesis have been heavily researched and understood as attractive therapeutic targets. Recently, evidence of their propensity for adding new sequences or domains during ARS evolution hints at broader functions and complexity outside of translation (42). Protein translation as a drug target has been validated for anti-infective compounds for a wide array of microbes (43). The natural product known as febrifugine, a quinazolinone alkaloid, and its analogues have shown antiparasitic activity in targeting ProRS. Halofuginone, a halogenated derivative of febrifugine, has shown promising potency though a lack of specificity, in that it inhibits both the parasite and human ProRS (43).

The structure of *Nf*ProRS folds into a α2 homodimer (Figure 6A) with each subunit containing three domains characteristic of Class II ARSs: the catalytic domain, the anticodon binding domain, and the editing domain (Figure 6B). The *Nf*ProRS catalytic domain features the three motifs which are exclusively conserved between class II ARSs for both sequence and structure-function (Figure 6C). Motif 1 is located at the interface of the dimer and is hypothesized to facilitate communication between the active sites of the two subunits (44). Motif 2 consists of β-strands connected by a variable loop which makes critical contacts with the acceptor stem of tRNA^Pro^ and thus plays an important role in proper tRNA recognition (45). Motif 3 is made up of entirely hydrophobic residues and comprises an integral part of the aminoacylation active site.

**Figure 6.**
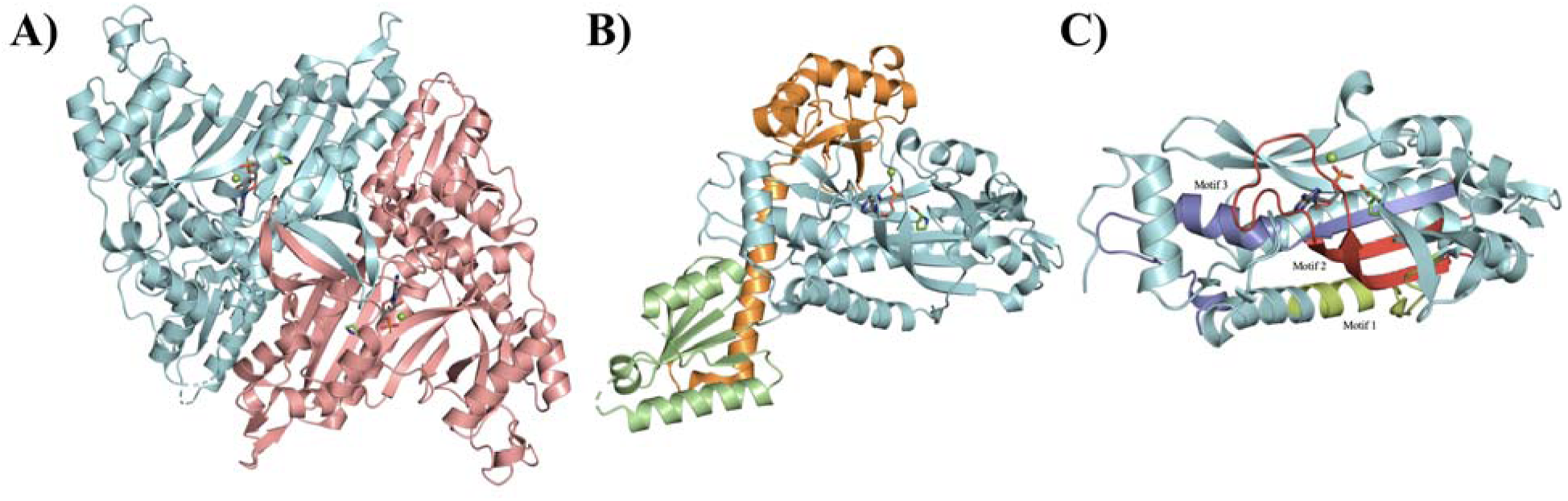
Structure of the *Nf*ProRS. (**A**) Both the biological and asymmetric unit of the structure are homodimeric. Individual polypeptide chains are shown in cyan and salmon. AMP, proline, and magnesium ligand molecules are also shown in yellow, purple, and green; respectively. (**B**) The three structure-function domains of *Nf*ProRS. The catalytic, anticodon binding, and editing domains are colored cyan, green, and orange; respectively. (**C**) The three highly conserved sequence motifs that characterize class II ARSs. Motif 1, colored lime, comprises the dimer interface. Motif 2, colored red, forms part of the acceptor stem. Motif 3, colored blue, is involved in forming the activated prolyl-adenylate.

Alignment of *Nf*ProRS bound to AMP and proline ligands (PDB: 6NAB) with apo *Hs*ProRS (PDB: 4K87) exhibits no significant structural changes between the apo and ligand forms of the ARS. The eukaryotic and archaeal origins of these ProRS make them suitable comparisons for the reason mentioned earlier: their strict conservation in all three structural domains. Both the proline and AMP bound *Nf*ProRS (PDB: 6NAB) and the halofuginone and AMP-PNP bound *Nf*ProRS (PDB: 6UYH) structures have been solved. The proline and AMP *Nf*ProRS (6NAB) shares structural homology with the halofuginone liganded ProRS (6UYH) and halofuginone binding induces a conformational change of residues 80-87 of the *N. fowleri* enzyme. In the proline bound 6NAB structure, residues 80-88 form a two-turn alpha helix (α4 in Figure 7). However, the halofuginone compound displaces Phe87 and disrupts the short helical structure.

**Figure 7.**
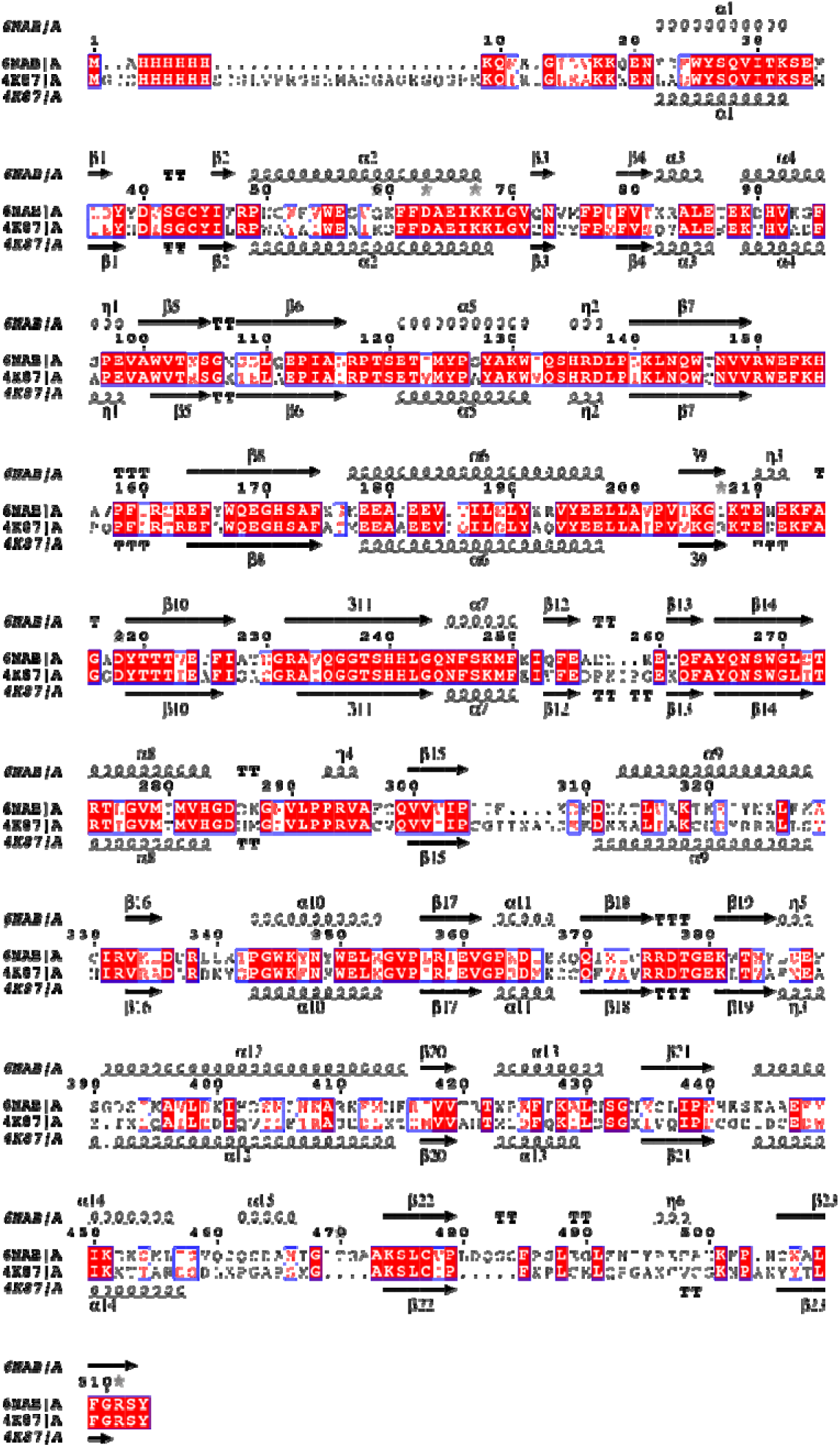
Alignment of *Nf*ProRS against its human homolog. Red background denotes residue conservation between the two structures, black text with white background signifies differences and red text encompassed in a blue box demonstrates differences in residues that are not categorized as significant and the residues belong to the same grouping. Secondary structure annotations signaling helices and sheet are reflective of both structures respectively. Other structures used in alignment; *H. sapiens* (PDB: 4K87_A).

Residues making up this helix (EKDHVEGFS) are disordered in the 6UYH coordinate set. The equivalent region of the human ProRS (PDB: 4K87) is structurally homologous to the proline bound *N. fowleri* in the absence of halofuginone binding. The equivalent helix in 4K87, residues 90-98 (EKTHVADFA), includes non-conservative amino acid substitution adjacent to the crucial phenylalanine which must be displaced for halofuginone to bind the human enzyme, including E85-G86 which are A-D in the human sequence. Exploiting differences in the mobility of this non-conserved loop adjacent to the active site of *Nf*ProRS and *Hs*ProRS could enable selective targeting. In addition, allosteric inhibitors that take advantage of sequence differences throughout the *Nf*ProRS might be found by screening, as was the case for *P. falciparum* ProRS (PfProRS) (46). Thus, ProRS may be a reasonable drug target for *N. fowleri* drug development.

## 3. CONCLUSION

This manuscript reports 19 new protein structures from *N fowleri* that are potential targets for structure-based drug discovery. Eighteen of 19 possess a >38% difference in AA alignment in comparison to the human homologs, suggesting selective inhibitors may be found by screening campaigns. In this paper we analyzed five of the *N. fowleri* enzymes that have ligands that define the active sites and compared them to human homologs. Though all are somewhat homologous at the active site, differences in four of the five *N. fowleri* enzymes analyzed support the hypothesis that selective active site inhibitors could be developed as therapeutics.

There are therapeutic opportunities, as well for some of the other 14 unexamined proteins as well. For example, the *Nf* serine tRNA synthetase (*Nf*SerRS) structure (PDB: 6BLJ). SerRS is required for charging tRNAs with serine critical for protein synthesis and thus is an essential gene. An insertion of four residues (391–395) adjacent to the substrate and tRNA binding sites creates a pocket with differential sequence identity to *Hs*SerRS and provides a foothold for the design of selective inhibitors blocking tRNA charging.

Even if selective active site inhibitors cannot be identified, high-throughput screening of compound libraries can still reveal selective inhibitors, as was found for *Plasmodium falciparum* ProRS compared with human *Hs*ProRS (46). In this case, two allosteric inhibitors were found to bury themselves into a lobe of the *Pf*ProRS enzyme, distant from the active site, and inhibit the activity of the *Pf*ProRS enzyme, but not *Hs*ProRS. Selective high throughput screening of a eukaryotic enzyme including counter screening against the homologous human enzyme, can also identify selective inhibitors as has been shown by us in the case of *Plasmodium* N- myristoyltransferase (32). Focusing on essential genes and drug targets of other eukaryotes and producing a pool of potential drug target structures, SSGCID has created a foundation on which to build structure-based drug discovery. The relatively quick successful progress through the pipeline has catalyzed a consortium of investigators interested in addressing *N. fowleri* drug discovery.

## 4. MATERIALS AND METHODS

### 4.1. Bioinformatics

The complete genome and transcriptome is available on the EupathDB BRC website (www.amoebadb.org) (2). The complete ORFs and annotated predicted proteome from *Naegleria fowleri* strain ATCC30863 was downloaded from AmoebaDB release 24. Analysis of the ORFs indicated that 39% were missing a start codon and 12% were missing a stop codon. The sequence authors, the Wittwer group at the Spiez Laboratory, confirmed that the 40% of transcripts without an AUG start codon were most likely due to the ORF finder they used, which searches for the longest ORFs in the RNA assembly, but has no start codon finding function. To address this issue, we applied a conservative strategy to select high quality sequences from the draft genome. A sequence homology search using BlastP against DrugBank v.4.3 targets (4,212 sequences) (15) and potential drug targets in the SSGCID pipeline (9,783 sequences) was performed. Sequences with at least 50% amino acid sequence identity over 70% coverage were selected for further filtering. Manual inspection indicated that half the potential targets without a start codon appeared to be significantly truncated when compared to the *Naegleria gruberi* and other closely related Eukaryota orthologues. Therefore, additional filters were applied to remove likely truncated sequences: (1) targets without a start or stop codon were discarded, (2) remaining candidates were blasted against the *Naegleria gruberi* proteome and sequences with over 10% length difference to their *Naegleria gruberi* orthologues were discarded, and (3) shorter variants with 100% match to a longer ORF transcript were discarded. In the end, 178 ORFs with a start and stop codon were identified, nominated, and approved by the SSGCID target selection board and NIAID to attempt structure determination.

### 4.2. High-throughput Protein Expression and Purification

All proteins discussed were PCR-amplified using cDNA as a template. RNA template of *Naegleria fowleri* ATCC30215 was provided by Dr. Christopher Rice (University of Georgia, Athens) through RNA extraction and cDNA synthesis using previously published methodology in *Acanthamoeba* (47). PCR, cloning, screening, sequencing, expression screening, large-scale expression and purification of proteins were performed as described in previous SSGCID publications (17, 48). All described constructs were cloned into a ligation-independent cloning (LIC) pET-14b derived, N-terminal His tag expression vector, pBG1861. Targets were expressed using chemically competent *E.coli* BL21(DE3)R3 Rosetta cells and grown in large-scale quantities in an auto-induction media (49). All purifications were performed on an AKTAexplorer (GE) using automated IMAC and SEC programs in adherence to prior established procedures (17).

### 4.3. Crystallization and Structure Determination

Crystal trials, diffraction, and structure solution were performed as previously published (16). Protein was diluted to 20 mg/mL and single crystals were obtained through vapor diffusion in sitting drops directly. The screens and conditions that yielded the crystals are listed in **Supplementary Table 1**. The screens that were used to find the crystallization conditions were typically JCSG+ (Rigaku Reagents), MCSG1 (Microlytic/Anatrace), Morpheus (Molecular Dimensions), in some cases supplemented by ProPlex (Molecular Dimensions) and JCSG Top96 (Rigaku Reagents). All data was integrated and scaled with *XDS* and *XSCALE* (50). Structures were solved by molecular replacement with *MOLREP* (51–53), *as implemented in MoRDa.* The structures were refined in iterative cycles of reciprocal space refinement in *Phenix* and real space refinement in *Coot* (54, 55). The quality of all structures was continuously checked using *MolProbity* (56) as implemented in Phenix. Structural comparisons for analysis among homologues was done using DALI Protein Structure Comparison Server.

## 5. ACKNOWLEDGMENTS

This work was funded by National Institute of Allergy and Infectious Diseases contract numbers HHSN272201700059C and HHSN272201200025C. This research used resources of the Advanced Photon Source, a U.S. Department of Energy (DOE) Office of Science User Facility operated for the DOE Office of Science by Argonne National Laboratory under Contract No. DE-AC02-06CH11357. Use of the LS-CAT Sector 21 was supported by the Michigan Economic Development Corporation and the Michigan Technology Tri-Corridor (Grant 085P1000817). We thank the SSGCID cloning and protein production groups at the Seattle Children’s Research Institute and at the University of Washington and Allison Pires for assistance with the crystallography table. We would like to thank Erin Egan, Veda Gadiraju, Caroline Francis, Becca Marks, and Matthew Howard who worked with Craig L. Smith on analyses of many of the *N. fowleri* reported structures, but their analyses were not reported in this paper.

