## Supplemental Table 1 for "*Naegleria fowleri:* protein structures to facilitate drug discovery for the deadly, pathogenic free-living amoeba"

**X-ray Crystallographic Data Collection**

| Description | malate dehydrogenase (mitochondrial) (MDH2) | s-adenosyl-L-homocysteine hydrolase | GDP-l-fucose synthetase | |
|---|---|---|---|---|
| # aminoacids | 436 | 472 | 333 | |
| Ligand | - | NAD, Adenosine | apo | NADP |
| PDB ID | 6UM4 | 5V96 | 6AQY | 6AQZ |
| Crystallization Condition | JCSG+ C4 | MCSG1 H11 | Morpheus C8 | MCSG1 H11 |
| Data Collection Statistics | | | | |
| Diffraction source | APS 21-ID-F | APS 21-ID-F | APS 21-ID-F | APS 21-ID-G |
| Wavelength (Å) | 0.97872 | 0.97872 | 0.97856 | 0.97856 |
| Temperature (K) | 100 K | 100 K | 100 K | 100 K |
| Detector | Rayonix MX-300 | Rayonix MX-300 | Rayonix MX-300 | Rayonix MX-300 |
| Crystal-detector distance (mm) | 200 | 250 | 325 | 320 |
| Rotation range per image (°) | 1 | 1 | 1 | 1 |
| Total rotation range (°) | 240 | 150 | 200 | 180 |
| Space group | P1 | P 21 21 21 | P 1 21 1 | P 1 21 1 |
| *a, b, c (Å)* | 60.33, 137.48, 139.28 | 69.83, 134.33, 239.79 | 99.73, 102.10, 121.82 | 97.37, 104.03, 120.17 |
| α, β, γ (°) | 91.093, 89.998, 91.482 | 90, 90, 90 | 90, 107.63, 90 | 90, 108.67, 90 |
| Mosaicity (°) | 0.176 | - | 0.238 | 0.239 |
| Resolution range (Å) | 44.23-2.05 (2.10-2.05) | 48.4-2.0 (2.05-2.00) | 35.6-2.55 (2.62-2.55) | 46.12-2.40 (2.46-2.40) |
| Total No. of reflections | 718,188 (52,953) | 953,738 (70,383) | 316,588 (23,761) | 331,976 (25,1030 |
| No. of unique reflections | 271,476 (19937) | 152,740 (11,197) | 75,854 (5557) | 87,872 (6513) |
| Completeness (%) | 96.7 (96.1) | 99.9 (100) | 99.7 (99.9) | 98.8 (98.8) |
| Redundancy | 2.65 (2.65) | 6.24 (6.29) | 4.17 (4.28) | 3.78 (3.84) |
| ⟨ I/σ(I)⟩ | 8.18 (2.0) | 16.01 (3.89) | 19.6 (2.7) | 14.96 (2.14) |
| *Rmeas* | 10.8 (65.0) | 10.1 (54.3) | 5.5 (64.6) | 5.9 (67.3) |
| *Rsym* | 8.5 (51.2) | 9.3 (49.8) | 4.8 (56.7) | 5.1 (57.9) |
| Overall B factor from Wilson plot (Å2) | 23.36 | 19.98 | 57.53 | 53.45 |
| Refinement Statistics | | | | |
| Resolution range (Å) | 44.23-2.05 (2.11-2.05) | 48.08-2.00 (2.05-2.00) | 35.6-2.55 (2.61-2.55) | 46.12-2.40 (2.46-2.40) |

| | | | | |
|---|---|---|---|---|
| Completeness (%) | 96.5 (96) | 99.92 (100) | 99.9 (100) | 98.82 (99) |
| σ cutoff | 1.98 | 1.34 | 1.34 | -3 |
| No. of reflections, working set | 270,670 | 152,736 (10,634) | 75,816 (4894) | 87,834 (5702) |
| No. of reflections, test set | 1853 (123) | 1977 (129) | 2059 (134) | 2041 (132) |
| Final Rcryst (%) | 21.06 (31.48) | 13.83 (16.27) | 18.57 (27.03) | 17.78 (25.75) |
| Final Rfree (%) | 23.24 (38.18) | 17.97 (20.69) | 24.03 (32.98) | 22.44 (31.15) |
| No. of non-H atoms | | | | |
| Protein | 28,960 | 14,187 | 14,496 | 14,847 |
| Solvent | 2,237 | 1,716 | 150 | 250 |
| Hetero | 16 | 484 | 22 | 244 |
| Total | 31,213 | 16,387 | 14,668 | 15,341 |
| R.m.s. deviations | | | | |
| Bonds (Å) | 0.007 | 0.007 | 0.008 | 0.008 |
| Angles (°) | 0.841 | 0.868 | 0.897 | 0.953 |
| Average B factors (Å2) | 33.15 | 22.53 | 71.42 | 67.02 |
| Ramachandran plot | | | | |
| Most favoured (%) | 99.3 | 97.3 | 98.05 | 97.92 |
| Allowed (%) | 0.7 | 2.7 | 1.95 | 1.92 |
| Outliers | 0 | 0 | 0 | 0.16 |
| Method | Molecular replacement | Molecular replacement | Molecular replacement | Molecular replacement |
| Starting model | 1BDM | 3OND | 4E5Y | 4E5Y |

| **Description** | **GDP-mannose-dehydratase** | **nucleoside diphosphate kinase** | **ubiquitin-conjugating enzyme e2** | |
|---|---|---|---|---|
| # aminoacids | 380 | 151 | 161 | |
| Ligand | NADP, GDP | - | | |
| PDB ID | 5UZH | 5U2I | 6MJ9 | 5V0R |
| Crystallization Condition | Morpheus F12 | MCSG1 G10 | Morpheus E9 | Top96 C9 |
| Data Collection Statistics | | | | |
| Diffraction source | APS 21-ID-F | APS 21-ID-F | Rigaku FR-E+ | Rigaku FR-E+ |
| Wavelength (Å) | 0.97872 | 0.97872 | 1.5418 | 1.5418 |
| Temperature (K) | 100 K | 100 K | 100K | 100K |
| Detector | Rayonix MX-300 | Rayonix MX-300 | Rigaku Saturn 944+ | Rigaku Saturn 944+ |
| Crystal-detector distance (mm) | 260 | 170 | 50 | 50 |

| | | | | |
|---|---|---|---|---|
| Rotation range per image (°) | 1 | 1 | 0.5 | 0.5 |
| Total rotation range (°) | 120 | 200 | 2x 180 | 3x 180 |
| Space group | P 42 21 2 | P 1 21 1 | P 21 21 2 | P 21 21 2 |
| *a, b, c (Å)* | 92.73, 92.73, 94.66 | 67.90, 110.38, 68.14 | 37.30, 42.87, 89.96 | 52.36, 85.24, 33.85 |
| α, β, γ (°) | 90, 90, 90 | 90, 118.55, 90 | 90, 90, 90 | 90, 90, 90 |
| Mosaicity (°) | 0.213 | 0.12 | 0.29 | 0.203 |
| Resolution range (Å) | 46.37-2.25 (2.31-2.25) | 40-1.40 (1.44-1.40) | 50-1.85 (1.90-1.85) | 42.62-1.55 (1.59-1.55) |
| Total No. of reflections | 193,943 (14,335) | 717,800 (48,952) | 125,625 (5516) | 244.943 (6428) |
| No. of unique reflections | 20,179 (1462) | 172,534 (12,732) | 12,875 (928) | 22,634 (1593) |
| Completeness (%) | 99.9 (100) | 99.6 (99.8) | 99.9 (99.9) | 99.7 (97.4) |
| Redundancy | 9.6 (9.8) | 4.16 (2.84) | 9.76 (5.94) | 10.8 (4.3) |
| ⟨ I/σ(I)⟩ | 24.67 (5.18) | 9.08 (3.37) | 30.4 (2.94) | 48.14 (6.26) |
| *Rmeas* | 7.9 (59.7) | 13.0 (46.80) | 4.7 (64.7) | 2.8 (22.7) |
| *Rsym* | 7.5 (56.6) | 11.3 (40.5) | 4.5 (58.9) | 2.7 (19.8) |
| Overall B factor from Wilson plot (Å2) | 33.65 | 8.2 | 31.29 | 15.26 |
| Refinement Statistics | | | | |
| Resolution range (Å) | 46.36-2.25 (2.30-2.25) | 40.57-1.40 (1.43-1.40) | 38.7-1.85 (1.92-1.85) | 42.62-1.55 |
| Completeness (%) | 99.9 (100) | 99.8 (100) | 99.9 (100) | 99.68 (97) |
| σ cutoff | 1.34 | 1.36 | 1.35 | 1.35 |
| No. of reflections, working set | 20,178 (1270) | 172,502 (10,557) | 12,875 (1242) | 22,634 (1295) |
| No. of reflections, test set | 2004 (150) | 2194 (172) | 1311 (138) | 2106 (131) |
| Final Rcryst (%) | 14.68 (17.01) | 14.10 (19.71) | 18.32 (27.14) | 15.61 (19.64) |
| Final Rfree (%) | 19.90 (24.19) | 16.89 (21.97) | 22.21 (29.76) | 19.10 (23.81) |
| No. of non-H atoms | | | | |
| Protein | 2,712 | 7,056 | 1,116 | 1,142 |
| Solvent | 159 | 1,523 | 114 | 246 |
| Hetero | 101 | 68 | 0 | 2 |
| Total | 2,972 | 8,647 | 1,230 | 1,390 |
| R.m.s. deviations | | | | |
| Bonds (Å) | 0.007 | 0.010 | 0.006 | 0.005 |
| Angles (°) | 0.88 | 1.11 | 0.849 | 0.889 |
| Average B factors (Å2) | 36.81 | 12.29 | 28.55 | 20.38 |
| Ramachandran plot | | | | |
| Most favoured (%) | 97.38 | 98.0 | 97.2 | 97.87 |

| | | | | |
|---|---|---|---|---|
| Allowed (%) | 2.62 | 2 | 2.8 | 2.13 |
| Outliers | 0 | 0 | 0 | 0 |
| Method | Molecular replacement | Molecular replacement | Molecular replacement | Molecular replacement |
| Starting model | 1T2A | 3VVT | 5V0R | 4R62 |

| **Description** | **glyceraldehyde-3-phosphate dehydrogenase** | **ras-related c3 botulinum toxin substrate 1 isoform x2** | **trafficking protein particle complex subunit 3** | **phosphoglycerate mutase** |
|---|---|---|---|---|
| # aminoacids | 333 | 200 | 187 | 250 |
| Ligand | NAD | | | |
| PDB ID | 6NLX | 5VCU | 6AQ3 | 5VVE |
| Crystallization Condition | MCSG1 H11 | MCSG1 B1 | JCSG+ H10 | MCSG1 A7 |
| Data Collection Statistics | | | | |
| Diffraction source | APS 21-ID-F | APS 21-ID-G | APS 21-ID-G | Rigaku FR-E+ |
| Wavelength (Å) | 0.97872 | 0.97856 | 0.97856 | 1.5418 |
| Temperature (K) | 100 K | 100 K | 100 K | 100K |
| Detector | Rayonix MX-300 | Rayonix MX-300 | Rayonix MX-300 | Rigaku Saturn 944+ |
| Crystal-detector distance (mm) | 210 | 235 | 295 | 50 |
| Rotation range per image (°) | 1 | 0.25 | 1 | 0.5 |
| Total rotation range (°) | 180 | 180 | 360 | 3x 180 |
| Space group | P 21 21 21 | P 65 2 2 | P 1 | I 2 2 2 |
| *a, b, c (Å)* | 73.01, 119.94, 162.90 | 48.38, 48.38, 624.56 | 53.64, 78.67, 80.57 | 77.61, 89.33, 157.88 |
| α, β, γ (°) | 90, 90, 90 | 90, 90, 120 | 114.55, 87.67, 95.02 | 90, 90, 90 |
| Mosaicity (°) | 0.208 | 0.096 | 0.245 | 0.25 |
| Resolution range (Å) | 49.67-1.80 (1.85-1.80) | 35.86-1.85 (1.90-1.85) | 33.38-2.40 (2.46-2.40) | 47.05-1.70 (1.74-1.70) |
| Total No. of reflections | 760,791 (60,276) | 789,781 (58,540) | 177,369 (13,408) | 751,758 (24,468) |
| No. of unique reflections | 129,448 (9616) | 39,544 (2800) | 45,852 (3373) | 60,168 (4118) |
| Completeness (%) | 97.4 (98.6) | 99.9 (100) | 98.1 (97.6) | 99.3 (93.1) |
| Redundancy | 5.88 (6.27) | 19.97 (20.91) | 3.87 (3.98) | 12.49 (5.94) |
| ⟨ I/σ(I)⟩ | 13.90 (2.88) | 26.17 (5.32) | 18.62 (2.75) | 34.11 (3.69) |
| *Rmeas* | 8.4 (60.0) | 7.8 (55.4) | 5.0 (63.6) | 4.4 (47.3) |
| *Rsym* | 7.6 (54.8) | 7.6. (54.0) | 4.3 (55.0) | 4.2 (43.1) |

| | | | | |
|---|---|---|---|---|
| Overall B factor from Wilson plot (Å2) | 27.54 | 27.55 | 53.96 | 19.42 |
| Refinement Statistics | | | | |
| Resolution range (Å) | 49.47-1.80 (1.85-1.80) | 35.87-1.85 (1.90-1.85) | 33.38-2.40 (2.46-2.40) | 47.05-1.70 (1.74-1.70) |
| Completeness (%) | 97.36 (99) | 99.81 (1000 | 98.22 (97) | 99.22 (92) |
| σ cutoff | 1.34 | 1.35 | 1.99 | 1.34 |
| No. of reflections, working set | 129,402 (9113) | 39,482 (2554) | 45,811 (3090) | 60,142 (3557) |
| No. of reflections, test set | 2109 (150) | 1979 (141) | 2024 (152) | 2059 (136) |
| Final Rcryst (%) | 16.85 (23.64) | 17.59 (23.39) | 17.12 (25.48) | 16.49 (22.42) |
| Final Rfree (%) | 20.39 (30.98) | 20.97 (31.38) | 20.86 (30.74) | 19.47 (29.09) |
| No. of non-H atoms | | | | |
| Protein | 10,175 | 2,835 | 7,715 | 3,965 |
| Solvent | 1,037 | 399 | 111 | 605 |
| Hetero | 192 | 58 | 146 | 29 |
| Total | 11,404 | 3,292 | 7,972 | 4,599 |
| R.m.s. deviations | | | | |
| Bonds (Å) | 0.006 | 0.006 | 0.008 | 0.008 |
| Angles (°) | 0.888 | 0.941 | 0.92 | 0.928 |
| Average B factors (Å2) | 29.47 | 35.78 | 67.44 | 27.72 |
| Ramachandran plot | | | | |
| Most favoured (%) | 96.16 | 97.54 | 99.2 | 97.94 |
| Allowed (%) | 3.54 | 2.46 | 0.8 | 2.06 |
| Outliers | 0.3 | 0 | 0 | 0 |
| Method | Molecular replacement | Molecular replacement | Molecular replacement | Molecular replacement |
| Starting model | 1U8F | 2QME | 1SZ7 | 1E58 |

| **Description** | **heterotrimeric G-protein alpha subunit Galpha7** | **Serine--tRNA ligase** | **dUTP pyrophosphatase** | |
|---|---|---|---|---|
| # aminoacids | 321 | 477 | 147 | |
| Ligand | | | UTP | apo |
| PDB ID | 6NE6 | 6BLJ | 6MJK | 5VJY |
| Crystallization Condition | Wizard 3/4 C8 | MCSG1 B8 | JCSG+ C6 | MCSG1 C12 |
| Data Collection Statistics | | | | |
| Diffraction source | APS 21-ID-F | APS 21-ID-G | APS 21-ID-G | APS 21-ID-G |
| Wavelength (Å) | 0.97872 | 0.97656 | 0.97856 | 0.97856 |

| | | | | |
|---|---|---|---|---|
| Temperature (K) | 100 K | 100 K | 100 K | 100 K |
| Detector | Rayonix MX-300 | Rayonix MX-300 | Rayonix MX-300 | Rayonix MX-300 |
| Crystal-detector distance (mm) | 235 | 245 | 175 | 240 |
| Rotation range per image (°) | 1 | 1 | 1 | 1 |
| Total rotation range (°) | 150 | 120 | 140 | 140 |
| Space group | P 21 21 21 | P 31 2 1 | P 21 3 | P 63 |
| *a, b, c (Å)* | 60.69, 68.31, 79.57 | 168.19, 168.19, 111.50 | 74.23, 74.23, 74.23 | 118.64, 118.64, 98.98 |
| α, β, γ (°) | 90, 90, 90 | 90, 90, 90 | 90, 90, 90 | 90, 90, 120 |
| Mosaicity (°) | 0.191 | 0.121 | 0.135 | 0.127 |
| Resolution range (Å) | 48.26-1.70 (1.74-1.70) | 42.05-2.10 (2.15-2.10) | 33.19-1.45 (1.49-1.40) | 34.25-2.00 (2.05-2.00) |
| Total No. of reflections | 222,267 (13,464) | 787,734 (58,505) | 416,266 (30,576) | 476,445 (35,320) |
| No. of unique reflections | 37,029 (2695) | 105,635 (7738) | 24,498 (1801) | 53,461 (3960) |
| Completeness (%) | 99.8 (99.8) | 99.8 (100.0) | 100 (100) | 100 (100) |
| Redundancy | 6.0 (4.9) | 7.45 (7.56) | 16.99 (16.97) | 8.91 (8.92) |
| ⟨ I/σ(I)⟩ | 23.09 (2.99) | 18.67 (4.31) | 46.35 (5.79) | 26.77 (4.12) |
| *Rmeas* | 5.0 (57.20 | 7.7 (58.4) | 4.1 (51.2) | 5.8 (56.5) |
| *Rsym* | 4.6 (51.2) | 7.1 (54.4) | 3.9 (49.7) | 5.4 (53.3) |
| Overall B factor from Wilson plot (Å2) | 19.7 | 31.28 | 21.78 | 30.44 |
| Refinement Statistics | | | | |
| Resolution range (Å) | 48.25-1.70 (1.74-1.70) | 42.047-2.10 (2.15-2.10) | 33.19-1.45 (1.52-1.45) | 34.28-2.00 (x) |
| Completeness (%) | 96.55 (88) | 99.79 (100) | 99.98 (100) | 99.96 (100) |
| σ cutoff | 0 | 1.36 | 1.35 | 1.35 |
| No. of reflections, working set | 35,817 (2033) | 105,584 (7320) | 24,472 (1510) | 53,440 (3398) |
| No. of reflections, test set | 2146 (96) | 1960 (147) | 2039 (129) | 2050 (139) |
| Final Rcryst (%) | 16.09 (22.50) | 15.82 (17.98) | 13.41 (12.92) | 14.23 (18.21) |
| Final Rfree (%) | 19.47 (24.85) | 19.11 (21.74) | 17.04 (18.68) | 17.65 (21.95) |
| No. of non-H atoms | | | | |
| Protein | 2,636 | 10,450 | 1,082 | 4,996 |
| Solvent | 334 | 1,029 | 166 | 498 |
| Hetero | 30 | 81 | 27 | 74 |
| Total | 3,000 | 11,560 | 1,275 | 5,568 |
| R.m.s. deviations | | | | |

| | | | | |
|---|---|---|---|---|
| Bonds (Å) | 0.008 | 0.008 | 0.005 | 0.006 |
| Angles (°) | 0.786 | 0.86 | 0.783 | 0.879 |
| Average B factors (Å2) | 24.13 | 40.25 | 20.64 | 39.59 |
| Ramachandran plot | | | | |
| Most favoured (%) | 96.39 | 98.87 | 98.59 | 98.61 |
| Allowed (%) | 3.28 | 1.13 | 1.41 | 1.39 |
| Outliers | 0.33 | 0 | 0 | 0 |
| Method | Molecular replacement | Molecular replacement | Molecular replacement | Molecular replacement |
| Starting model | 5DO9 | 3LSQ | 4OOP | 4OOP |

| **Description** | **rab GDP dissociation inhibitor alpha** | **protein arginine N-methyltransferase** | | **polyubiquitin with 3 ub domains** |
|---|---|---|---|---|
| # aminoacids | 444 | 328 | | 230 |
| Ligand | | apo | SAH | |
| PDB ID | 6C87 | 6CU3 | 6CU5 | 5VIX |
| Crystallization Condition | JCSG+ E1 | JCSG+ A3 | ProPlex F12 | 25% PEG 3350, 0.2M AmSO4, 0.1M Na Cacodylate, pH 5.5 |
| Data Collection Statistics | | | | |
| Diffraction source | APS 21-ID-F | APS 21-ID-F | APS 21-ID-F | APS 21-ID-F |
| Wavelength (Å) | 0.97872 | 0.97872 | 0.97872 | 1 |
| Temperature (K) | 100 K | 100 K | 100 K | 100 K |
| Detector | Rayonix MX-300 | Rayonix MX-300 | Rayonix MX-300 | Rayonix MX-300 |
| Crystal-detector distance (mm) | 345 | 325 | 300 | 200 |
| Rotation range per image (°) | 1 | 1 | 1 | 1 |
| Total rotation range (°) | 150 | 130 | 200 | 200 |
| Space group | P 21 21 21 | P 21 21 21 | P 1 21 1 | P 1 21 1 |
| *a, b, c (Å)* | 97.34, 103.12, 195.55 | 249.63, 101.46, 105.89 | 43.20, 101.45, 80.68 | 34.02 67.71 35.83 |
| α, β, γ (°) | 90, 90, 90 | 90, 90, 90 | 90, 100.10, 90 | 90, 114.49, 90 |
| Mosaicity (°) | 0.134 | 0.112 | 0.303 | 0.157 |
| Resolution range (Å) | 47.95-2.60 (2.67-2.60) | 46.99-2.50 (2.56-2.50) | 42.75-2.70 (2.77-2.70) | 50-1.55 (1.59-1.55) |
| Total No. of reflections | 380,965 (28.200) | 504,864 (37,484) | 80,274 (5837) | 88,835 (6282) |

| | | | | |
|---|---|---|---|---|
| No. of unique reflections | 61,169 (4471) | 93675 (6834) | 18861 (1356) | 21,353 (1549) |
| Completeness (%) | 99.8 (100) | 99.9 (100) | 99.9 (1000 | 99.2 (99.7) |
| Redundancy | 6.23 (6.31) | 5.3 (5.49) | 4.25 (4.30) | 4.16 (4.06) |
| ⟨ I/σ(I)⟩ | 19.54 (3.42) | 16.18 (3.11) | 9.41 (2.79) | 14.00 (2.31) |
| *Rmeas* | 8.1 (59.4) | 8.5 (61.6) | 13.4 (64.0) | 6.1 (65.3) |
| *Rsym* | 7.5 (54.5) | 7.7 (55.8) | 11.7 (56.1) | 5.4 (56.7) |
| Overall B factor from Wilson plot (Å2) | 45.51 | 40.86 | 38.64 | 19.21 |
| Refinement Statistics | | | | |
| Resolution range (Å) | 47.95-2.60 (2.67-2.60) | 46.99-2.50 (x) | 42.75 (x) | 32.6-1.55 (1.59-1.55) |
| Completeness (%) | 99.75 (100) | 99.84 (100) | 99.71 (100) | 99.4 (100) |
| σ cutoff | 1.34 | 1.35 | 1.34 | 1.36 |
| No. of reflections, working set | 61,129 (4174) | 93,624 (6029) | 18,806 (1266) | 21,330 (1862) |
| No. of reflections, test set | 1937 (131) | 2166 (132) | 1868 (143) | 1862 (152) |
| Final Rcryst (%) | 17.19 (24.30) | 19.1 (21.91) | 19.6 (25.10) | 17.09 (25.78) |
| Final Rfree (%) | 22.25 (33.49) | 23.3 (26.08) | 26.6 (37.41) | 20.41 (32.19) |
| No. of non-H atoms | | | | |
| Protein | 13,406 | 14,035 | 4,908 | 1,291 |
| Solvent | 396 | 458 | 73 | 170 |
| Hetero | 10 | 72 | 52 | 0 |
| Total | 13,812 | 14,565 | 5,033 | 1,461 |
| R.m.s. deviations | | | | |
| Bonds (Å) | 0.005 | 0.008 | 0.008 | 0.007 |
| Angles (°) | 0.711 | 0.936 | 1.026 | 0.937 |
| Average B factors (Å2) | 52.39 | 54.88 | 45.88 | 31.25 |
| Ramachandran plot | | | | |
| Most favoured (%) | 98.35 | 97.83 | 97.27 | 99.34 |
| Allowed (%) | 1.65 | 2.11 | 2.73 | 0.66 |
| Outliers | 0 | 0.06 | 0 | 0 |
| Method | Molecular replacement | Molecular replacement | Molecular replacement | Molecular replacement |
| Starting model | 3CPI | 3Q7E | 6CU3 | 5H07 |

| **Description** | **peptidylprolyl isomerase** | **prolyl-tRNA synthetase** |
|---|---|---|
| # aminoacids | 119 | 535 |

| Ligand | | | Pro/AMP | Halofuginone/AMpPNP |
|---|---|---|---|---|
| PDB ID | 6MKE | 6B4P | 6NAB | 6UYH |
| Crystallization Condition | MCSG1 E12 | Morpheus H11 | TOP96 D7 | MSCG1 A6 |
| Data Collection Statistics | | | | |
| Diffraction source | Rigaku FR-E+ | APS 21-ID-F | APS 21-ID-G | APS 21-ID-G |
| Wavelength (Å) | 1.5418 | 0.97872 | 0.97656 | 0.97656 |
| Temperature (K) | 100K | 100 K | 100 K | 100 K |
| Detector | Rigaku Saturn 944+ | Rayonix MX-300 | Rayonix MX-300 | Rayonix MX-300 |
| Crystal-detector distance (mm) | 50 | 200 | 295 | x |
| Rotation range per image (°) | 0.5 | 1 | 1 | 1 |
| Total rotation range (°) | 3x 360 | 200 | 180 | 200 |
| Space group | P 1 21 1 | P 1 21 1 | C 1 2 1 | C 1 2 1 |
| *a, b, c (Å)* | 59.05, 60.16, 73.34 | 37.25, 61.15, 45.76 | 156.93, 66.10, 112.98 | 157.29, 66.54, 112.56 |
| α, β, γ (°) | 90, 97.12, 90 | 90, 90.28, 90 | 90, 102.03, 90 | 90, 101.74, 90 |
| Mosaicity (°) | 0.142 | 0.125 | 0.117 | 0.169 |
| Resolution range (Å) | 48.683-2.05 (2.10-2.05) | 45.76-1.55 (1.59-1.55) | 38.37-2.00 (2.05-2.00) | 50.0-1.75 (1.80-1.75) |
| Total No. of reflections | 524,516 (28,257) | 123,738 (8923) | 279,947 (15,053) | 478,997 (35,437) |
| No. of unique reflections | 31313 (2306) | 29,144 (2123) | 76,263 (5333) | 114,792 (8431) |
| Completeness (%) | 97.1 (96.2) | 97.6 (97.2) | 99.4 (94.4) | 99.9 (99.9) |
| Redundancy | 16.75 (12.25) | 4.25 (4.20) | 3.67 (2.82) | 4.17 (4.20) |
| ⟨ I/σ(I)⟩ | 37.0 (8.41) | 16.32 (5.45) | 12.78 (2.24) | 15.69 (2.52) |
| *Rmeas* | 6.2 (27.0) | 6.1 (24.5) | 9.4 (60.6) | 6.0 (59.2) |
| *Rsym* | 6.0 (25.9) | 5.3 (21.4) | 8.0 (49.5) | 5.2 (51.7) |
| Overall B factor from Wilson plot (Å2) | 30.287 | 17.06 | 29.58 | 22.34 |
| Refinement Statistics | | | | |
| Resolution range (Å) | 48.67 - 2.05 (2.10-2.05) | 45.76-1.55 (1.59-1.55) | 38.0 -2.00 (2.05-2.00) | 42.66-1.75 (1.79-1.75) |
| Completeness (%) | 97.13 (96) | 97.6 (97) | 99.4 (94) | 99.9 (100) |
| σ cutoff | 1.35 | 1.39 | 1.35 | 1.35 |
| No. of reflections, working set | 31,308 (2095) | 29,135 (1889) | 76,253 (4973) | 114,783 (7973) |
| No. of reflections, test set | 1965 (118) | 2024 (1460 | 1991 (143) | 1918 (154) |
| Final Rcryst (%) | 21.34 (23.25) | 15.47 (18.15) | 15.12 (22.67) | 15.49 (22.28) |

| | | | | |
|---|---|---|---|---|
| Final Rfree (%) | 25.67 (28.69) | 18.67 (22.68) | 19.16 (26.53) | 18.07 (23.73) |
| No. of non-H atoms | | < | | |
| Protein | 3,600 | 1,764 | 7,919 | 7,889 |
| Solvent | 388 | 222 | 802 | 888 |
| Hetero | 228 | 0 | 116 | 206 |
| Total | 4,216 | 1,986 | 8,837 | 8,983 |
| R.m.s. deviations | | | | |
| Bonds (Å) | 0.002 | 0.005 | 0.007 | 0.006 |
| Angles (°) | 0.53 | 0.742 | 0.843 | 0.831 |
| Average B factors (Å2) | 29.21 | 24.68 | 29.48 | 30.45 |
| Ramachandran plot | | | | |
| Most favoured (%) | 98.04 | 98.23 | 98.07 | 97.95 |
| Allowed (%) | 1.96 | 1.77 | 1.93 | 1.84 |
| Outliers | 0 | 0 | 0 | 0.21 |
| Method | Molecular replacement | Molecular replacement | Molecular replacement | Molecular replacement |
| Starting model | 6B4P | 1R9H | 4HVC | 6NAB |
